# CRISPR/Cas12a mediated genome engineering in photosynthetic bacteria

**DOI:** 10.1101/2020.10.05.327569

**Authors:** Yang Zhang, Jifeng Yuan

## Abstract

Purple non-sulfur photosynthetic bacteria (PNSB) such as *R. capsulatus* serve as a versatile platform for fundamental studies and various biotechnological applications. In this study, we sought to develop the class II RNA-guided CRISPR/Cas12a system from *Francisella novicida* for both genome editing and gene down-regulation in *R. capsulatus*. About 90% editing efficiency was achieved by using CRISPR/Cas12a driven by a strong promoter P_*puc*_ when targeting *ccoO* or *nifH* gene. When both genes were simultaneously targeted, the multiplex gene editing efficiency reached >63%. In addition, CRISPR interference using deactivated Cas12a was also evaluated using reporter genes *gfp* and *lacZ*, and the repression efficiency reached >80%. In summary, our work represents the first report to develop CRISPR/Cas12a mediated genome editing/transcriptional repression in *R. capsulatus*, which would greatly accelerate PNSB-related researches.

**IMPORTANCE:** Purple non-sulfur photosynthetic bacteria (PNSB) such as *R. capsulatus* serve as a versatile platform for fundamental studies and various biotechnological applications. However, lack of efficient gene editing tools remains a main obstacle for progressing in PNSB-related researches. Here, we developed CRISPR/Cas12a for genome editing via the non-homologous end joining (NHEJ) repair machinery in *R. capsulatus*. In addition, DNase-deactivated Cas12a was found to simultaneously suppress multiple targeted genes. Taken together, our work offers a new set of tools for efficient genome engineering in PNSB such as *R. capsulatus*.

## INTRODUCTION

Purple non-sulfur photosynthetic bacteria (PNSB) such as *Rhodobacter capsulatus* are facultative anaerobic Gram-negative bacteria and are generally regarded as safe (GRAS) (1, 2). These bacteria have the versatile metabolic ability to grow under a variety of habitats (3). For instance, *R. capsulatus* could use a wide spectrum of carbon sources including short-chain fatty acids, dicarboxylic acids, sugars, agricultural and food wastes for chemotrophic growth under aerobic to microaerobic conditions (4). It can also utilize the organic or inorganic compounds (such as S_2_O_3_^2-^, H_2_S, and Fe^2+^) as electron donors for photoheterotrophic growth with energy from light to assimilate CO_2_ under anaerobic condition (5, 6). Moreover, when the nitrogen source is scarce, *R. capsulatus* can perform nitrogen fixation and produce hydrogen via the action of nitrogenases (7). However, due to the lack of photosynthetic system II, the hydrogen donor for photosynthesis is organic or inorganic substrates instead of water, and no oxygen is released during the process (8).

To survive under such diverse living conditions, PNSB has evolved a complex metabolic network with highly specialized enzyme complexes and regulatory mechanisms. Recently, PNSB has attracted special attention as a platform microorganism for biotechnological applications such as H_2_ production as an alternative renewable energy to fossil fuels (7, 9) and polyhydroxybutyrate as biodegradable plastics (10, 11), bioremediation of wastewater (12) and heavy metal (13), fixation of CO_2_ and N_2_ (14, 15), expression of membrane proteins (16, 17) and metalloenzymes (18, 19). More importantly, as the synthesis of photopigments in PNSB relies on isoprenoid biosynthetic pathway, it offers a robust and effective isoprenoid metabolism. Therefore, PNSB has been regarded as a promising host for isoprenoid biosynthesis (1, 2, 20).

Currently, the synthetic biology tools for engineering PNSB are lacking, which significantly slows the progress in PNSB-related biotechnological applications. The genetic tools in PNSB still heavily rely on traditional homologous recombination using suicide plasmids (9), gene transfer agent (GTA) derived from a phage-like particle capable of transduction (21), and gene mutation using transposon (22). There is a pressing need to develop new and efficient genetic tools to accelerate the researches in PNSB. Recently, Clustered Regularly Interspaced Short Palindromic Repeats (CRISPR) and CRISPR-associated protein (Cas) systems (23) have been developed as versatile tools for genome editing and transcription regulation in a variety of organisms (24, 25). The class II type II CRISPR/Cas9 system containing a Cas nuclease (Cas9), a trans-activating CRISPR RNA (tracrRNA), and a *cis*-regulatory RNA (crRNA) has been engineered as a powerful genetic tool in various organisms including PNSB such as *Rhodobacter sphaeroides* (26, 27).

Cas12a, a class II type V-A endonuclease, is characterized as a dual nuclease referring to crRNA processing, target-site recognition, and DNA cleavage (28). The CRISPR/Cas12a system also has been repurposed as a genetic tool adapted for many species (29, 30). Compared to CRISPR/Cas9 that recognizes the target locus close to a guanine-rich protospacer adjacent motif (PAM) sequence (NGG) to create blunt ends (26), Cas12a assisted by a mature crRNA binds the protospacer segment flanked by a thymidine-rich PAM sequence (TTN) to form staggered ends (28, 31), which might promote the non-homologous end joining (NHEJ) machinery to fix the double strand breaks (DSB). In addition, the compact design of crRNA and self-processing crRNA by Cas12a make CRISPR/Cas12a a more versatile and robust tool than CRISPR/Cas9 system. To the best of our knowledge, there is no report on CRISPR/Cas12a mediated genetic tools in PNSB. In this study, we aimed to investigate the feasibility of using CRISPR/Cas12a assisted genetic engineering tools in *R. capsulatus*. In addition, we also attempted to evaluate CRISPR/dCas12a as artificial transcriptional factors for targeted gene knock-down applications.

## RESULTS

### Evaluation of CRISPR/Cas12a mediated genome editing in *R. capsulatus*

To begin with, we first evaluated the potential toxicity of Cas12a from *Francisella novicida* expression in *R. capsulatus*. The *E. coli*-*Rhodobacter* shuttle vector pBBRdMCS, a multicopy vector derived from pBBR1MCS2 (32), was used to express the *sacB* expression cassette and *cas12a* gene under the control of P_*lac*_ promoter, resulting the plasmid pBdRCas12a (Figure 1). Notably, P_*lac*_ promoter has constitutive transcription activity due to the absence of the *lacI* repressor gene in *R. capsulatus* genome (33). When the plasmid pBdRCas12a in parallel with the control plasmid pBdRSacB were transformed into *R. capsulatus*, the conjugation efficiency for pBdRCas12a was close to that achieved by the control (Figure. 2a). Considering the size of pBdRCas12a (10876 bp) plasmid was doubled as compared to the control (5144 bp), it was likely that the slight decrease in conjugation efficiency could occur. Therefore, these findings suggested that Cas12a is non-toxic to *R. capsulatus* under the tested conditions.

**Figure 1.**
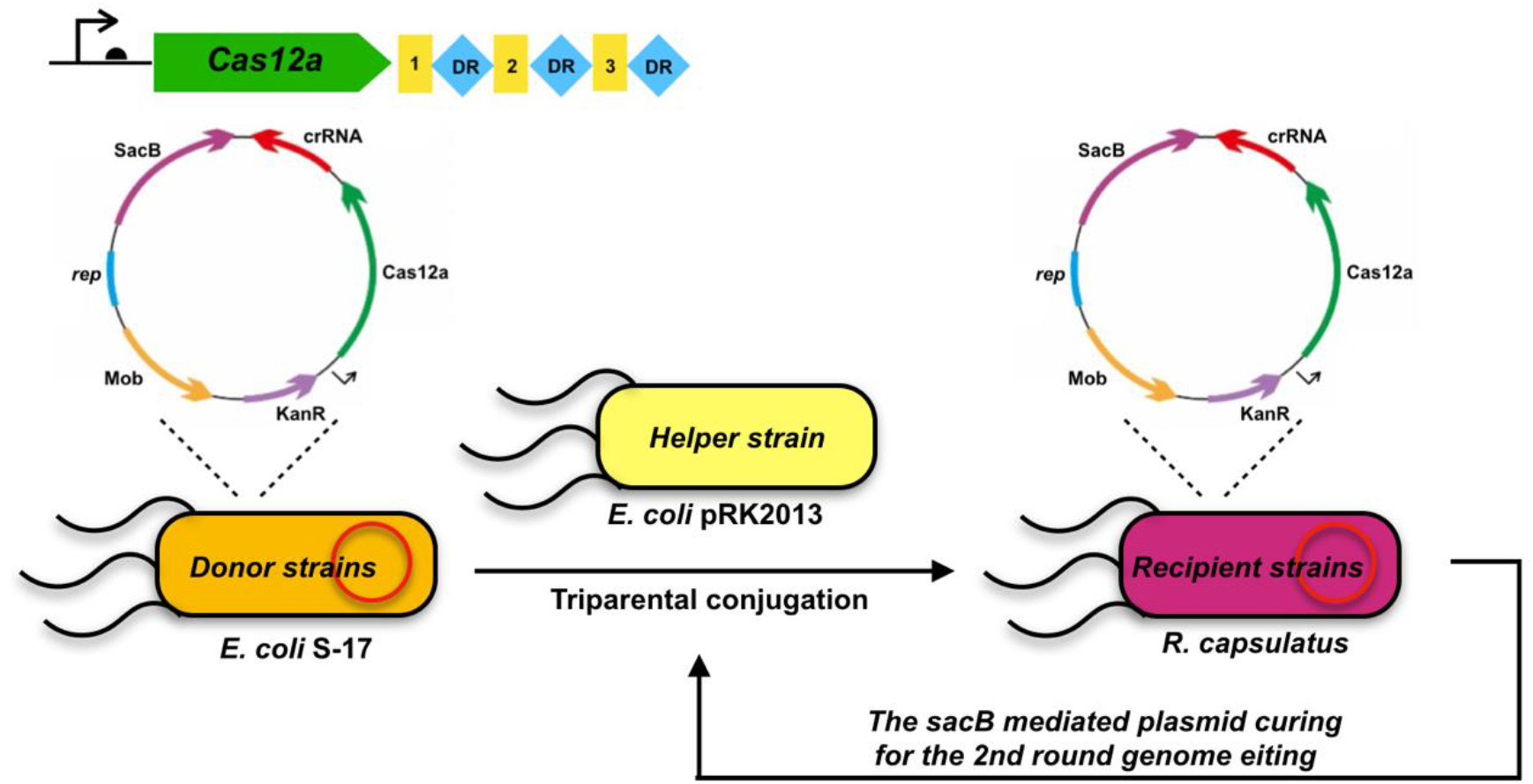
Schematic diagram of CRISPR-Cas12a system for genome editing in *R. capsulatus*. Cas12a and crRNA were expressed from a single plasmid with kanamycin marker, shuttle replicon, and mobilization for transferring plasmid from donor strains *E. coli* into recipient strains of *R. capsulatus*. The *sacB* cassette was used as the counter-selection marker for curing plasmid. CRISPR/Cas12a system was first constructed in *E. coli* S-17, then conjugated into *R. capsulatus* via the helper strain of *E. coli* pRK2013.

**Figure 2.**
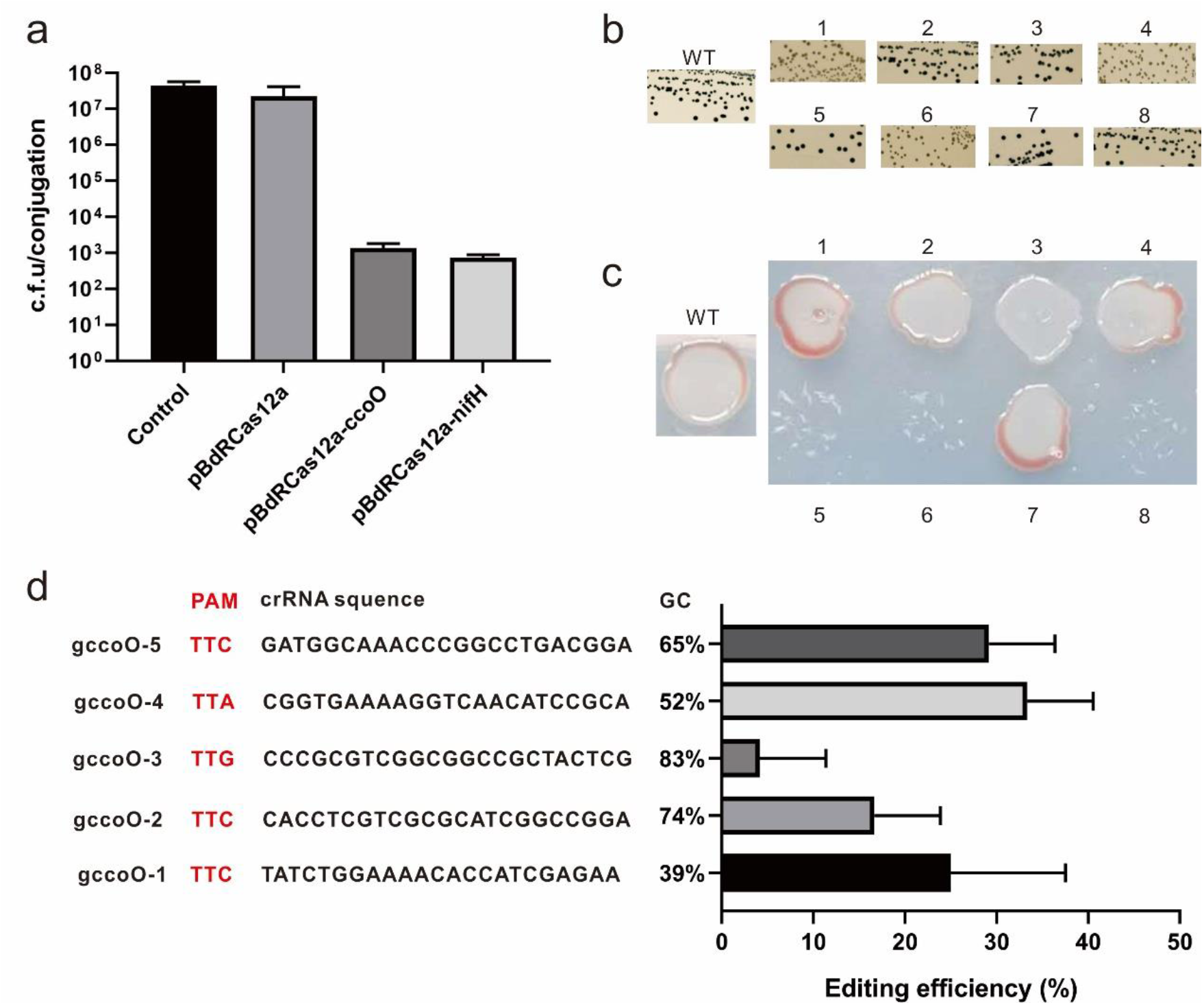
Genome editing using CRISPR-Cas12a in *R. capsulatus*. (a) Colony forming unit (c.f.u) of *R. capsulatus* conjugation mixtures containing Cas12a with or without the expression of crRNA target. (b) Verification of *ccoO* disruption by Nadi staining. (c) Verification of *nifH* disruption by growing on the MedA agar plate without nitrogen source. (d) Comparison of the genome editing efficiency of five crRNAs with different GC contents. Experiments were carried out in triplicate, and the data are presented as mean ± S.D.

To confirm whether Cas12a is functional in *R. capsulatus, ccoO* gene encoding a subunit of *cbb*_3_-cytochrome oxidase (*cbb*_3_-Cox) and *nifH* gene encoding a subunit of nitrogenase were chosen as the disruption targets. The *in vivo cbb*_*3*_-Cox activity of *R. capsulatus* strains could be visualized using Nadi staining solution. As nitrogenase is essential for the nitrogen fixation, thus *nifH* mutants would abolish the growth under nitrogen source depleting conditions. According to the literatures, crRNA to achieve the sufficient efficiency for genome editing typically comprises a 19-nt direct repeat (DR) and a 23-nt guide sequence (28). Therefore, the crRNA-expressing module was assembled into the pBdRCas12a plasmid under the control of the same promoter as Cas12a. Based on this principle, plasmid pBdRCas12a-ccoO and pBdRCas12a-nifH were constructed to target genes of *ccoO* and *nifH*, respectively. As shown in Figure 2a, the conjugation efficiencies declined more than 4 orders of magnitude compared to the control plasmid. Since the generation of DSB at the targeted genes by CRISPR/Cas12a typically results in decreased survival rates (27), these findings suggested that the CRISPR/Cas12a system might be functional in *R. capsulatus*. As shown in Figure 2b and 2c, *ccoO* or *nifH* was successfully mutated by Cas12a mediated by NHEJ mechanism. However, the editing efficiency mediated by CRISPR/Cas12a was only 30%∼40%, which is not satisfactory when compared to the traditional methods. It was reported that the genome editing efficiency assisted by CRISPR/Cas system was observably impacted by GC content of guide RNA and the target positions (34). In this study, five crRNAs with different GC contents ranging from 39% to 83% were designed for *ccoO* (gccoO-1 to gccoO-5). As shown in Figure 2d, the crRNAs with different GC contents resulted in editing efficiencies ranging from 4% to 33%. In particular, similar efficiencies (25%, 33%, and 30%, respectively) were achieved for GC contents ranging from 39% to 65%, indicating that Cas12a was not as sensitive to the GC content as the Cas9 counterpart. However, higher GC content might affect the crRNA folding and maturity, thereby decreasing the genome editing.

### Optimization of CRISPR/Cas12a mediated genome editing in *R. capsulatus*

According to previous reports, efficient expression of Cas12a is crucial to achieve efficient CRISPR/Cas12a assisted genome editing systems (35). As the heterologous promoter P_*lac*_ is a relatively weak promoter, it might limit the CRISPR/Cas12a expression and result in the low editing efficiency. To search alternative strong promoters to drive the expression of CRISPR/Cas12a in *R. capsulatus*, four endogenous promoters including promoters from *puc* operon (encoding light-harvesting complex II), *puf* operon (encoding light-harvesting complex I), *pgk* gene (encoding phosphoglycerate kinase), and *eno* gene (encoding enolase) were chosen. Encouragingly, all promoter candidates exhibited much higher activity than P_*lac*_ (Figure 3a). Among the tested promoters, P_*puc*_ showed the highest activity, which corresponds to approximately 5.4-fold improvement over that of P_*lac*_.

**Figure 3.**
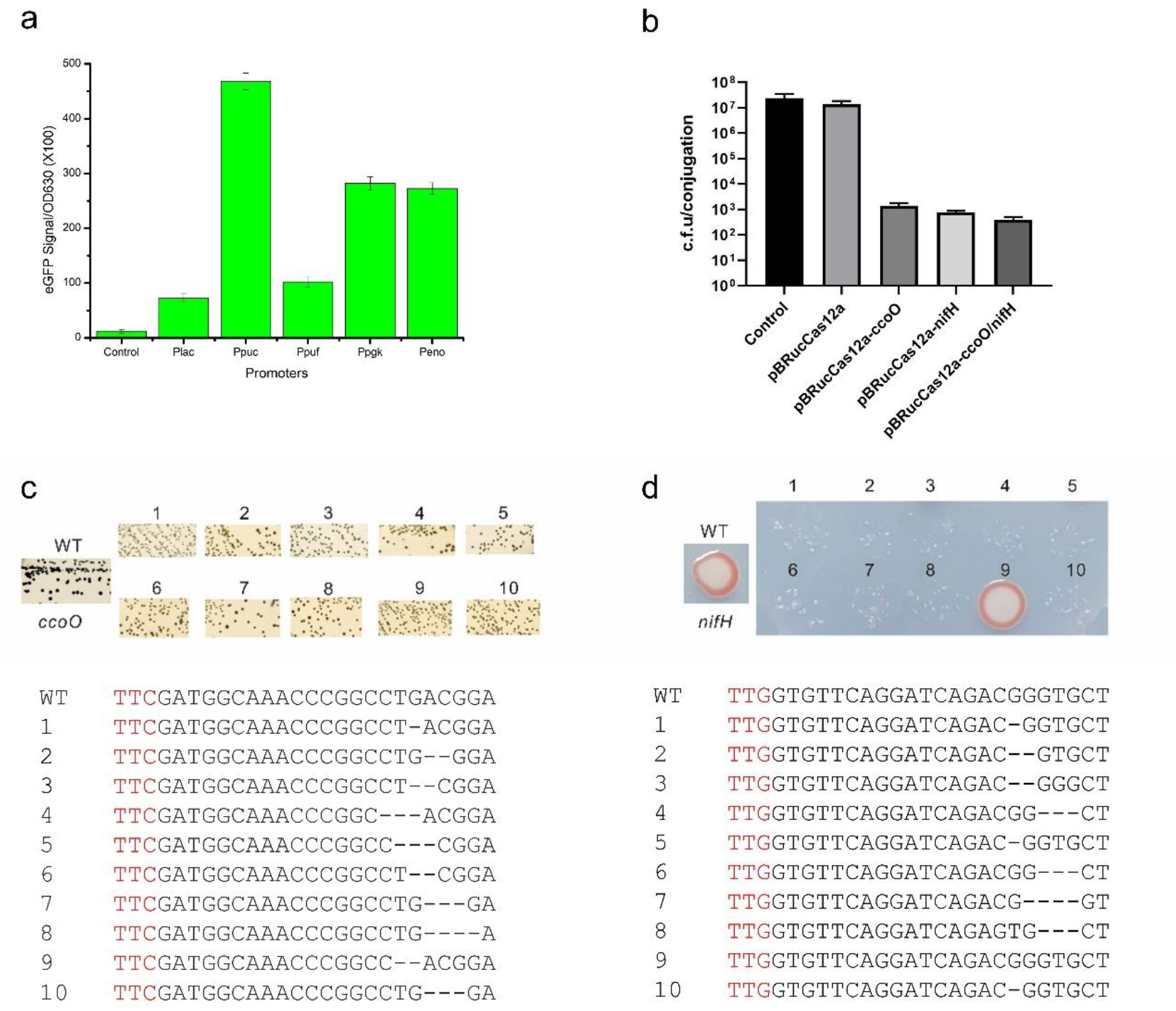
Optimization of CRISPR-Cas12a mediated genome editing in *R. capsulatus*. (a) Transcriptional activities of five promoters in *R. capsulatus* (P_*lac*_, P_*puc*_, P_*puf*_, P_*pgk*_, and P_*eno*_) as measured by eGFP fluorescence intensities. (b) Colony forming unit (c.f.u) obtained after conjugation of empty vector and Cas12a with or without the crRNA targets driven by promoter P_*puc*_. (c) Confirmation of *ccoO* disruption by Nadi staining and sequencing. (d) Confirmation of *nifH* disruption by growth without nitrogen source and sequencing. WT represents the wild type strain. Experiments were carried out in triplicate, and the data are presented as mean ± S.D.

On the basis of above findings, the promoter P_*puc*_ was applied to drive the expression of CRISPR/Cas12a system in the remaining studies. When compared to the first-generation design, the new versions of pBRucCas12a-ccoO and pBRucCas12a-nifH only differed in the promoter sequences (P_*puc*_ v.s. P_*lac*_). These plasmids were transformed into *R. capsulatus* with pBRPpucSacB as the control plasmid. As depicted in Figure 3b, the conjugation efficiency of Cas12a expression driven by P_*puc*_ was close to that of Cas12a driven by P_*lac*_, suggesting that higher expression of Cas12a was not toxic to *R. capsulatus*. As shown in Table 1, 10 randomly selected transformants on *ccoO* disruption was verified by Nadi staining and sequencing, and 93% editing efficiency was achieved. The editing efficiency for the disruption of *nifH* reached up to 87%. And it was observed that short indels (1 to 4 bp) occurred at the protospacer regions (Figure 3c and 3d). Noteworthily, the editing efficiency using CRISPR/Cas12a driven by a stronger promoter P_*puc*_ was much higher than that under the control of promoter P_*lac*_, suggesting that CRISPR/Cas12a could be an effective genome editing tool in PNSB such as *R. capsulatus*. In addition, approximately 63% positive transformants were observed during simultaneous disruption of *ccoO* and *nifH* as shown in Table 1.

**Table 1.** Editing efficiency using CRISPR/Cas12a driven by P_*puc*_ promoter.

| Target gene | Replicate | Editing efficiency | Average editing efficiency |
| --- | --- | --- | --- |
| *ccoO* | 1 | 100% (10/10) | 93% |
|  | 2 | 90% (8/10) |  |
|  | 3 | 90% (9/10) |  |
| *nifH* | 1 | 80% (7/10) | 87% |
|  | 2 | 90% (9/10) |  |
|  | 3 | 90% (8/10) |  |
| *ccoO, nifH* | 1 | 70% (7/10) | 63% |
|  | 2 | 60% (6/10) |  |
|  | 3 | 60% (6/10) |  |

### Transcriptional repression by CRISPR interference (CRISPRi) using deactivated Cas12a (dCas12a)

As evidenced by CRISPR/Cas12a mediated genome editing, it is possible to further employ CRISPR/dCas12a as an artificial transcriptional factor for targeted gene knockdown. The nuclease activity in the RuvC domain of Cas12a was abolished by introducing the E1006A mutation (36). The dCas12a driven by the promoter P_*puc*_ was cloned into a low copy *E. coli*-*Rhodobacter* shuttle vector pRK415 (37), to give plasmid pRKucdCas12a (Figure 4a). The transcriptional repression efficiency of CRISPRi was evaluated by employing the *gfp* and *lacZ* as the report genes, which were preintegrated at the *phbC* and *pucBA* sites, respectively. For each reporter gene, three crRNAs targeting at the template strand were designed (g1, g2, and g3 or z1, z2, and z3, as shown in Figure 4b), and one crRNA targeting at the non-template strand was examined (g4 or z4, Figure 4b). The above plasmids harboring dCas12a with different crRNAs were then transformed into *R. capsulatus* with chromosomal integrated expression cassettes of *gfp* and *lacZ*. As shown in Figure 4c and 4d, all the crRNAs targeting the template strand could lead to different degrees of repression of the *gfp* or *lacZ* expression. Similar to previous findings that crRNA-binding position near the translation initiation site on the template strand is more effective for CRISPRi (38, 39), we found the most efficient repression occurred when the crRNAs were designed near the start codon (*gfp* and *lacZ* repression, 84% and 82%, respectively). However, only ∼20% repression was achieved with targeting at the non-template strand for both reporter genes. To test the capability of multiplex gene regulation, dual crRNA expressing plasmid was constructed to simultaneously repress *gfp* and *lacZ* expressions. The expression levels of *gfp* and *lacZ* were simultaneously reduced by 80% and 77%, respectively. These findings indicated that the CRISPRi system based on dCas12a has broad utilities to implement multiplex transcriptional repressions in *R. capsulatus* for the future metabolic engineering applications.

**Figure 4.**
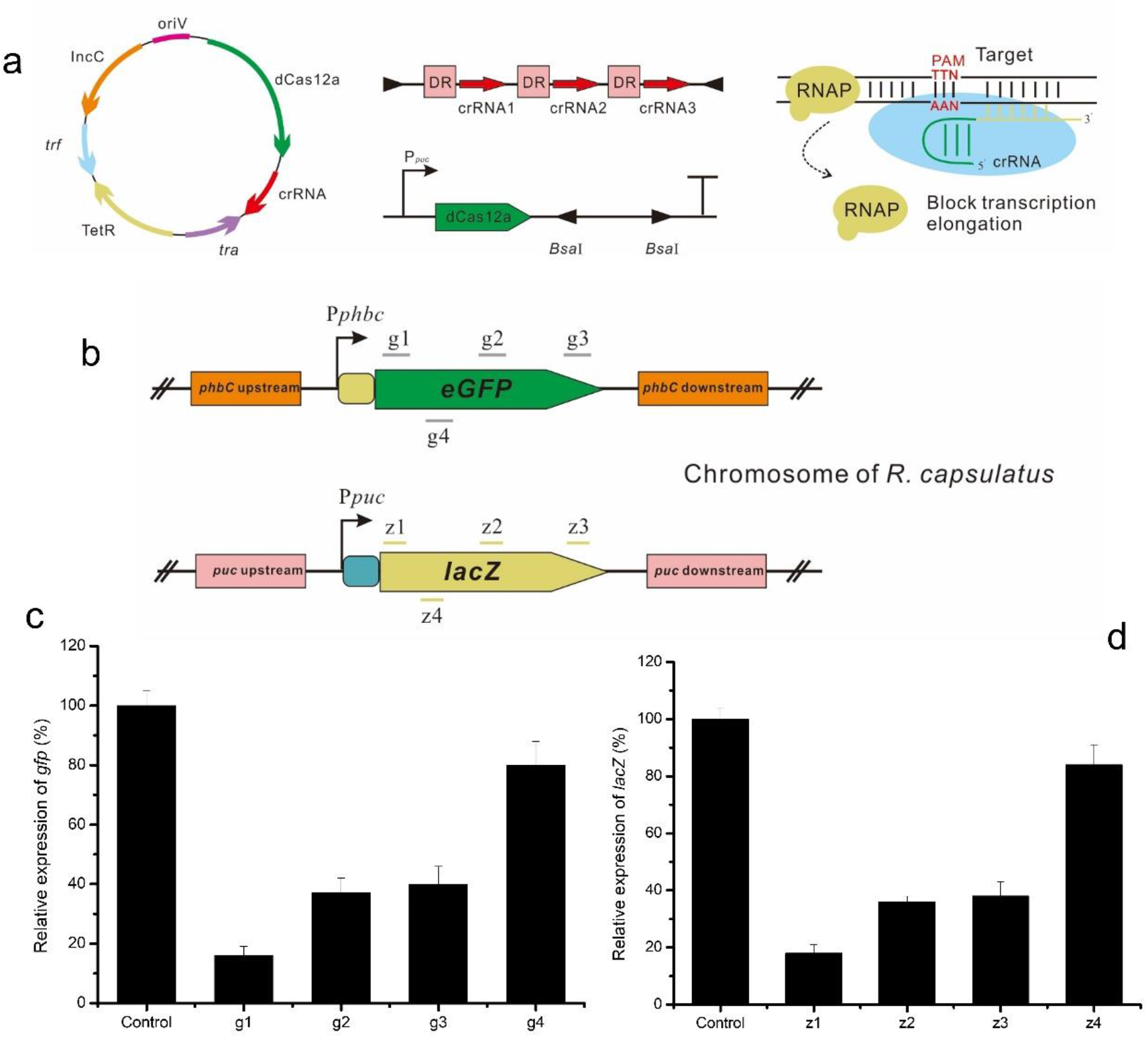
Transcriptional repression of *gfp* and *lacZ* by CRISPRi based on dCas12a. (a) dCas12a and crRNA driven by promoter P_*puc*_ were expressed from a single plasmid with tetracycline marker, shuttle replicon IncC, *tra* combined with *trf* for transferring plasmid into the host. (b) crRNAs targeting different positions on the template strand (g1, g2, and g3 or z1, z2, and z3) and the non-template strand (g4 or z4) of *gfp* or *lacZ* gene. (c) Comparison of *gfp* expression level suppressed by dCas12a when targeting at different positions of the template and non-template strand. (d) Comparison of *lacZ* expression level suppressed by dCas12a when targeting different positions of the template and non-template strand. Experiments were carried out in triplicate, and the data are presented as mean ± S.D.

## DISCUSSION

The CRISPR/Cas system is revolutionizing the field of genome editing in various organisms. The type V-A CRISPR/Cas12a system has the ability to self-process crRNA and cleave DNA to give staggered ends instead of blunt ends. Due to these characteristics, CRISPR/Cas12a assisted genome editing is a superior alternative to CRISPR/Cas9 (28). In this work, we developed an efficient genetic tool using Cas12a from *F. novicida* in *R. capsulatus*, a promising platform microorganism for PNSB- related biotechnological applications. Based on previous findings, Cas9 from *S. pyogenes* (SpCas9) is toxic at high expression in several bacteria such as *Corynebacterium glutamicum* (40) and *Cyanobacteria sp*. (41), which limits the application of CRISPR/Cas9 in these microorganisms. In this work, we found that Cas12a expression driven by strong promoters in a multicopy vector did not show noticeable toxicity to *R. capsulatus* like *C. glutamicum*.

Genome editing with CRISPR/Cas is typically based on DSB repaired by homologous recombination (HR) or NHEJ. As CRISPR/Cas12a creates a staggered end resulting a 5-nt 5’ overhang rather than a blunt cleavage product, which might promote the DSB repair mediated by NHEJ mechanism. Therefore, CRISPR/Cas12a was applied for genome editing via the NHEJ machinery in *R. capsulatus*. The initial studies revealed that about 30%∼40% editing efficiency was obtained when targeting *ccoO* or *nifH* gene based on NHEJ pathway, which was not satisfactory when compared to NHEJ- mediated CRISPR/Cas in *Streptomyces coelicolor* (42) and *E. coli* (43). To further improve the genome editing efficiency in *R. capsulatus*, a strong promoter of pigment operon *puc* (P_*puc*_) was identified to drive the CRISPR/Cas12a system, and ∼90% editing efficiency was achieved for single gene disruption, whereas 63% editing efficiency was obtained for two gene disruptions. Therefore, the CRISPR/Cas12a system based on the NHEJ mechanism has great potentials for the future multiplex genome editing. To the best of our knowledge, CRISPRi using DNase-deactivated Cas9 (dCas9) or dCas12a has not been reported in *Rhodobacter* species. Previous findings suggested that dCas12a exhibits higher repression efficiency with crRNAs targeting at the template strand over non-template strand, and the closer crRNA near to the translational start sites, the higher repression efficiency is obtained (35, 38, 44). In this study, similar results were also observed in *R. capsulatus* using dCas12a driven by P_*puc*_ when targeting at *gfp* or *lacZ* gene. More importantly, dual repression of *gfp* and *lacZ* genes reached a similar efficiency to that of single target, indicating that CRISPR/dCas12a might be applicable for multiplex transcription regulation in *R. capsulatus*.

## MATERIALS AND METHODS

### Strains and culture conditions

All plasmids were introduced by heat-shock into competent cells of *Escherichia coli* S17-1 or Top 10 cultured with Luria-Bertani (LB) medium containing appropriate antibiotics at 37°C. The wild type of *R. capsulatus* SB1003 was employed for genetic modifications and the strain was cultured in Sistrom’s mineral media (MedA) (45) at 35°C. Antibiotics were used at the following final concentrations: gentamycin 12 mg/mL, tetracycline 12.5 mg/L, kanamycin 50 mg/L for *E. coli*; gentamycin 12 mg/mL, tetracycline 2.5 mg/L, kanamycin 10 mg/L for *R. capsulatus*. All strains used in this study are listed in Supplementary Table S1.

### Plasmid constructions

All oligonucleotides utilized in this work are listed in Supplementary Table S2. Plasmids and genomic DNA were extracted by Biospin kits (Bioer Technology). PCR amplification was conducted by High Fidelity Phusion DNA polymerase or Taq polymerase from New England Biolab. Plasmids were constructed by standard restriction endonucleases and ligation approach followed by digestion analysis and DNA sequencing verification. The detailed procedures were described in Supplementary Materials and Methods. All the plasmid constructs utilized in this study are listed in Supplementary Table S3.

### Triparental conjugation procedure

Plasmids were transformed into *R. capsulatus* by triparental conjugation with the helper plasmid of pRK2013 (46). Before the conjugation, *R. capsulatus* was grown with 10 mL MedA at 35°C for 48 h. The *R. capsulatus* cells were centrifuged, collected and washed once with fresh MedA, and resuspended with 400 μL MedA. Meanwhile, *E. coli* HB101 harboring the helper plasmid pRK2013 and *E. coli* S17-1 carrying the desired plasmid were incubated in 5 mL LB medium supplemented with proper antibiotics at 37°C for overnight. The *E. coli* cells were centrifuged, collected and washed once with fresh MedA, and resuspended with 1 mL MedA, respectively. *R. capsulatus* of 100 μL as a recipient strain, *E. coli* HB101/pRK2013 of 30 μL as a helper strain and *E. coli* S17-1 of 30 μL as a donor strain were mixed and spotted into a MedA agar plate as a ∼2 cm diameter spot incubating at 35°C for 24 h. The mixed cells were centrifuged, collected, washed with fresh MedA and resuspended with 1 mL MedA. The resuspended cell culture of 100 μL was spread on a MedA agar plate with appropriate antibiotics and incubated at 35°C for 2-3 days until colony appeared.

### CRISPR/Cas12a mediated genome editing and transcriptional repression in *R. capsulatus*

For CRISPR/Cas12a assisted genome editing, pBdRCas12a or pBRucCas12a derivatives carrying the corresponding crRNAs were transformed into *R. capsulatus* by triparental conjugation for disrupting the target genes. For CRISPR/dCas12a mediated transcriptional repression, *gfp* and *lacZ* expression cassette were integrated at *phbC* and *pucBA* sites of the chromosome via the traditional method mediated by pZJD29c or pK18mobsacB as previously described (7). Briefly, several recombinants containing pZJD29c or pK18mobsacB derived plasmids were cultivated in MedA medium for overnight, and then spread on the MedA agar plate containing 10% sucrose. The colonies were then subjected to diagnostic PCR analysis to confirm the integration of *gfp* and *lacZ* cassette into the *phbC* and *pucBA* sites. Next, pRKucdCas12a derived plasmids carrying the corresponding crRNAs were transformed into *R. capsulatus* and selected on the MedA agar plate supplemented with tetracycline. The fluorescence intensities and LacZ activities were measured for comparing the transcriptional repression.

### *In vivo cbb*_*3*_-cytochrome oxidase activity

To test whether the *ccoO* gene encoding for a subunit of *cbb*_*3*_-cytochrome oxidase (*cbb*_*3*_-Cox) was disrupted by CRISPR/Cas12a system, the *in vivo cbb*_*3*_-Cox activity of *R. capsulatus* was visualized qualitatively using Nadi staining solution containing 35 mM α-naphthol and 30 mM N,N,N′,N′-dimethyl-p-phenylene diamine (DMPD) dissolved in 1:1 (vol/vol) ethanol and water (47). When the *ccoO* gene was successfully disrupted, the strains showed the natural color (red) due to inactivation of *cbb*_*3*_-Cox, otherwise the colonies appeared blue.

### Fluorescence intensities and *β*-galactosidase activity assays

To measure the eGFP fluorescence, *R. capsulatus* recombinants were cultured in 5 mL MedA medium for 48 h. Optical density at 630 nm (*OD*_630_) and fluorescence intensities were monitored by a microplate reader (Synergy H1, BioTek). The excitation of eGFP was set at 485 nm as well as emission at 510 nm. Fluorescence intensities were normalized to culture OD_630_ for comparing the relative expression level of eGFP. The *β*-galactosidase activity assay was performed as previously described (48). In brief, the active protein exacted from *R. capsulatus* reacted with o-nitrophenyl-β-D- galactopyranoside (ONPG) at 35°C. When the reaction is over, the mixture was employed to detect *β*-galactosidase activity using the following equation:

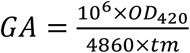

where *GA* is the *β*-galactosidase activity (μM ONPG-Hydrolyzed/(min ·mg-protein)), *OD*_420_ is the optical density at 420 nm, *t* is the reaction time (min) and *m* is the mass of the protein (mg).

## Author contributions

J.Y. conceived the project. Y.Z performed the experiments and collected the data. J.Y. and Y.Z. interpreted the data and wrote the manuscript.

## Acknowledgements

This work was supported by Xiamen University under grant no. 0660-X2123310, XMU Training Program of Innovation and Entrepreneurship for Undergraduates no. 2020Y1000 and 202010384194, and ZhenSheng Biotech, China.

## Competing financial interests

The authors declare no competing financial interests.

## Supporting Information

Supplementary materials and methods, list of strains used in the study (Table S1), list of oligonucleotides used in the study (Table S2) and list of plasmids used in the study (Table S3).

